# Measurement Quality Metrics to Improve Absolute Microbial Cell Counting

**DOI:** 10.1101/2025.02.18.638308

**Authors:** K. Parratt, D. Newton, J. Dunkers, J. Dootz, M. Hunter, A. Logan-Jackson, L. Pierce, S. Sarkar, S. Servetas, NJ. Lin

**Affiliations:** National Institute of Standards and Technology (NIST), Biosystems and Biomaterials Division, Gaithersburg, MD, USA; National Institute of Standards and Technology (NIST), Statistical Engineering Division, Boulder, CO, USA

**Keywords:** Measurement quality metrics, Absolute cell count, Microbial cell viability, Fluorescence flow cytometry, Impedance flow cytometry, CFU, Proportionality

## Abstract

Total and viable microbial cell counts are increasingly important for applications including live biotherapeutic products, food safety, and probiotics. In microbiology, cells are quantified using methods such as colony forming unit (CFU), flow cytometry, and polymerase chain reaction (PCR), but different methods measure different aspects of the cells (measurands), and results may not be directly comparable across methods. In the absence of a ground-truth reference material for cell count, one cannot quantify the accuracy of any cell counting method, which limits method performance assessments and comparisons. Herein, a modified analysis of cell counting methods based on the ISO 20391-2:2019 standard was developed and demonstrated for microbial cell samples diluted over a log-scale range of concentrations. *Escherichia coli* samples ranging in concentration from approximately 5 × 10^5^ cells/mL to 2 × 10^7^ cells/mL were quantified using CFU, Coulter principle, fluorescence flow cytometry, and impedance flow cytometry. Quality metrics modified from the ISO standard were calculated for each method and shown to be repeatable across replicate experiments. The quality metrics illustrate large differences in proportionality and variability across methods, with total cell counts in good agreement and viable cell count having more variability. As the ISO standard is meant to guide fit-for-purpose method selection, interpretation of the results and quality metrics can drive method choice and optimization. The framework introduced here will help researchers select fit-for-purpose counting methods for quantification of microbial total and viable cells across a range of applications.

## Introduction

High confidence total and viable microbial cell count measurements are challenging but critical for the rigorous characterization of microbial cell products, a growing field of biotechnology that includes live biotherapeutic products, probiotics, and microbial reference materials (Jackson et al., 2019, FDA-CFSAN, 2018, FDA-CBER, 2016a, FDA-CBER, 2016b). Quantifying the number of microbes present in a sample is also critical for other fields including food safety, water quality, antimicrobial testing, and microbiome characterization. Total microbial cell count is used as a normalization factor when reporting results (e.g., results “per cell”) and is a quantification of all microbes, live and dead, present in a sample. For example, there is great interest in optimizing molecular methods such as polymerase chain reaction (PCR) or genomic sequencing to accurately characterize microbial samples. However, results from these methods are challenging to interpret without a total cell count or genome count before DNA extraction to quantify measurement biases (Galazzo et al., 2020, McLaren et al., 2019). Viable microbial cell count can indicate the health of a microbial culture or product and is often used for potency measurements, such as for fecal microbiota transplantations (Ferring Pharmaceuticals Inc., 2022). Viable count is also important for microbial reference materials used to quantify limits of detection for sterility tests that support the safety and purity of biomanufactured products, such as cell and gene therapies (Gebo and Lau, 2020, Lin-Gibson et al., 2021).

Colony forming unit (CFU) assays are the bedrock of traditional microbiology and industrial microbial cell counts (Martini et al., 2024, Jasson et al., 2010, Clausen et al., 2018). CFU assays quantify culturable subpopulations based on the number of colonies that grow on solid media and represent a straightforward, accessible, and time-proven method often used synonymously with viable cell count. However, limitations in the CFU method, for example a long time-to-result or inability to count dead cells, drive industry toward alternative or complementary cell counting methods. Newer measurement technologies that report total and/or viable cell counts include fluorescence flow cytometry and impedance-based instruments. These methods can be quicker and higher throughput than CFU, but they do not explicitly measure cell growth. Instead, these methods quantify on other measurands, meaning the quantity intending to be measured (VIM, 2017). Fluorescence flow cytometry relies on scattered and fluorescent light to calculate total particle concentration and can also characterize cells using fluorescent probes. Many fluorescent probes are designed to measure cell health properties such as membrane integrity, membrane potential, metabolic activity, and DNA replication, though implementation typically requires optimization for each microbial sample of interest (Müller and Nebe-von-Caron, 2010, Veal et al., 2000). Impedance techniques can measure each particle passing through an aperture or channel and calculate the number of particles per unit volume. The BactoBox (SBT Instruments) and Multisizer (Coulter counter, Beckman Coulter) are two instruments^1^ that detect particles as changes in impedance though the detection technologies differ (Graham, 2022, Clausen et al., 2018, Bertelsen et al., 2023). Comparison among these measurands and understanding where differences arise can be critical drivers of protocol development for specific use cases for microbial cell counting (Jordal et al., 2023).

These different measurands may not agree for a sample of interest and each stakeholder will have unique cell counting needs, so many factors must be considered when selecting a cell counting method(s) for a particular application. Measurement quality varies with method and can often indicate that one method is more fit-for-purpose than another. While many stakeholders are interested in accuracy of a count value, accuracy cannot be determined without a certified cell-based reference material, and the material would need to be certified with values appropriate for each measurand of interest. For example, this might require a material certified for CFU, total cell count, and membrane potential. In the absence of such materials, other quality metrics must be relied upon to guide method selection. ISO 20391-2:2019: *Biotechnology – Cell counting – Part 2: Experimental design and statistical analysis to quantify counting method performance* provides a method to evaluate to what extent a cell counting method is proportional by measuring a stock solution diluted across a range of cell concentrations and calculating measurement quality metrics including proportionality, coefficient of variation, and R^2^ value (ISO, 2019).

Proportionality is a characteristic of an ideal measurement process whereby dilutions of a sample by a given factor should result in corresponding reductions in the measured values by the same factor and intersect the origin. For example, a sample diluted by half should result in a measurement that is half the concentration of the original sample. Previous work at NIST has demonstrated how the ISO standard can be applied to mammalian cells to compare counting methods (Sarkar et al., 2017, Pierce et al., 2023). The current work makes one large modification to the ISO 20391-2:2019 experimental protocol by allowing dilution factors that span more than one order of magnitude (i.e., are on the log scale). This modified design better accommodates the wide range of cell concentrations common in microbial experimental designs (e.g., antimicrobial effectiveness) or samples (e.g., natural microbiomes).

Here, the objective was to develop a modified ISO experimental design and analysis method suitable for logarithmic scale and demonstrate its application to compare microbial counting techniques. We measured a model laboratory species, *Escherichia coli NIST0056*, using four methods (CFU, Coulter principle, fluorescence flow cytometry, and impedance flow cytometry) for a total of six reported measurements (three total cell count measurements and three viable cell count measurements). Blinded samples that ranged in concentration from approximately 5 × 10^5^ cells/mL to 2 × 10^7^ cells/mL were quantified with each method operating with fixed acquisition conditions. A polymeric bead sample was also quantified with total particle count methods based on fluorescence and Coulter principle. To account for the fact that the dilution factors in this study were evenly spaced on a log-scale, the ISO standard calculations and statistical procedures were modified accordingly. Similar to the ISO standard, this work compares the proportionality, magnitude, and variability of cell count measurements across different counting methods. Additional analyses were also performed, including evaluating repeatability of metrics across multiple dates, quantifying different sources of experimental variability via hierarchical Bayesian modeling, determining the extent to which proportional “sub-ranges” exist for the various methods, and evaluating the impact of sample order. Results for this use case are discussed with a focus on how to determine which methods are fit-for-purpose. The demonstrated experimental design and mathematical analyses can be directly implemented by other microbial researchers who wish to characterize their methods to better quantify their microbes of interest.

## Materials and Methods

### Overview

The methods used to count the bacterial cells were CFU, Coulter principle (Graham, 2022), fluorescence flow cytometry, and impedance flow cytometry. The operator for each method, who also performed the data analysis, was held constant. From the three methods, six count datasets for cells were reported (**Table 1**). Raw data and analysis code are available at https://doi.org/10.18434/mds2-3410 (Parratt et al., 2025).

**Table 1:** List of Datasets and Methods. The instrument, measurand, underlying measurement principle, and method-specific dilutions are listed for each dataset.

| Dataset | Method Principle | Measurand | Instrument |
| --- | --- | --- | --- |
| Total-A and Bead-A | Coulter principle | Particle count | Multisizer 4 |
| Total-B | Impedance | Particle count | BactoBox |
| Total-C and Bead-C | Fluorescence | Particle count | CytoFLEX LX |
| Viable-B | Impedance | Intact count | BactoBox |
| Viable-C | Fluorescence | Non-intact count | CytoFLEX LX |
| Viable-D | Growth | Colony count (CFU) | Growth at 37°C |

Total particle counts were reported in the Multisizer, BactoBox, and CytoFLEX datasets. Viable cell counts were reported in the BactoBox, CytoFLEX, and CFU datasets. Experiments were performed on four separate dates with two biological replicates for CFU and Multisizer and three biological replicates for BactoBox and CytoFLEX (**Table 2**). Polymer bead counting was also performed on a single date using two of the instruments (CytoFLEX, Multisizer). It is critical to note that the quality metrics calculated here are a result of the *entire* measurement process, which includes the starting material, experimental operators, experimental processing steps, instrument acquisition settings, and data analysis. Thus, shorthand names are used for the datasets to avoid emphasizing method identities, since results cannot be extended to others’ measurement processes.

**Table 2:** Instrument List. List of instruments used for each Biological or Bead replicate experiment (Bio. Rep. with *E. coli* NIST0056 or Bead Rep.).

| Instrument | Bio. Rep. 1 | Bio. Rep. 2 | Bio. Rep. 3 | Bio. Rep. 4 | Bead Rep. 1 |
| --- | --- | --- | --- | --- | --- |
| BactoBox | Y | Y |  | Y |  |
| CFU* |  | Y |  | Y |  |
| CytoFLEX LX | Y | Y | Y |  | Y |
| Multisizer 4 |  | Y | Y |  | Y |

\*CFU was performed manually, not by an instrument.

### Cell Preparation

Study design was established to determine the range of cell concentrations, number of levels, and dilution factors to be tested. These determinations were specific to the instruments and cell type under consideration and should be modified as needed for other systems. Lyophilized bacterial cells were selected as the test sample, as many microbial reference materials are produced in a lyophilized format. Previous characterization of the lyophilized material (not shown) indicated that tube-to-tube variability was relatively low and that materials were stable at the recommended storage temperature of 4 °C. Four separate experiments were performed on different dates and are referred to as “biological replicates,” with a new cell stock made for each experiment. To generate a biological replicate, four lyophilized bacterial pellets (*Escherichia coli* NIST0056, custom manufactured by Microbiologics, Inc., stored at 4 °C) were equilibrated at room temperature and each rehydrated in 1 mL phosphate buffered saline (PBS, Corning, catalogue # 46-013-CM). All four rehydrated pellets were then added to one tube containing 36 mL PBS to generate a single sample of approximately 2 × 10^7^ cells/mL (**Figure 1**). This stock was then diluted into six dilution factors evenly spaced in log-scale, with three replicate samples (“sample replicates”) per dilution factor prepared separately for a total of 18 samples (**Figure 1, Table 3**). Samples were then divided into 4 aliquots and transferred to each operator with blinded labeling. Samples were provided in semi-random order with sample replicates for each dilution factor spaced throughout the experiment duration. Samples were stored at 4 °C until acquisition, for up to 5 h depending on the method and sample order. Each operator collected a pre-specified 1 or 2 technical replicate measurements of each sample (“observation replicates”) for a total of 5 observations per dilution factor or 30 acquisitions per method, as shown in **Figure 1**. Each operator reported final results before the samples were unblinded. This process was repeated for each biological replicate.

**Table 3:** Cell Concentrations. Based on the cell concentration of the cell stock solution, target dilution factors were expected to result in samples with the listed cell concentrations. The cell concentrations are calculated by multiplying the stock cell concentration by the dilution factor. All cell concentrations are approximate because no ground-truth reference material exists.

| Dilution Factor | 1 | 0.478 | 0.2285 | 0.1095 | 0.0525 | 0.025 |
| --- | --- | --- | --- | --- | --- | --- |
| Approximate Cell Concentration (cell/mL) | $2 \times 10^7$ | $9.56 \times 10^6$ | $4.57 \times 10^6$ | $2.19 \times 10^6$ | $1.05 \times 10^6$ | $5 \times 10^5$ |

**Figure 1:**
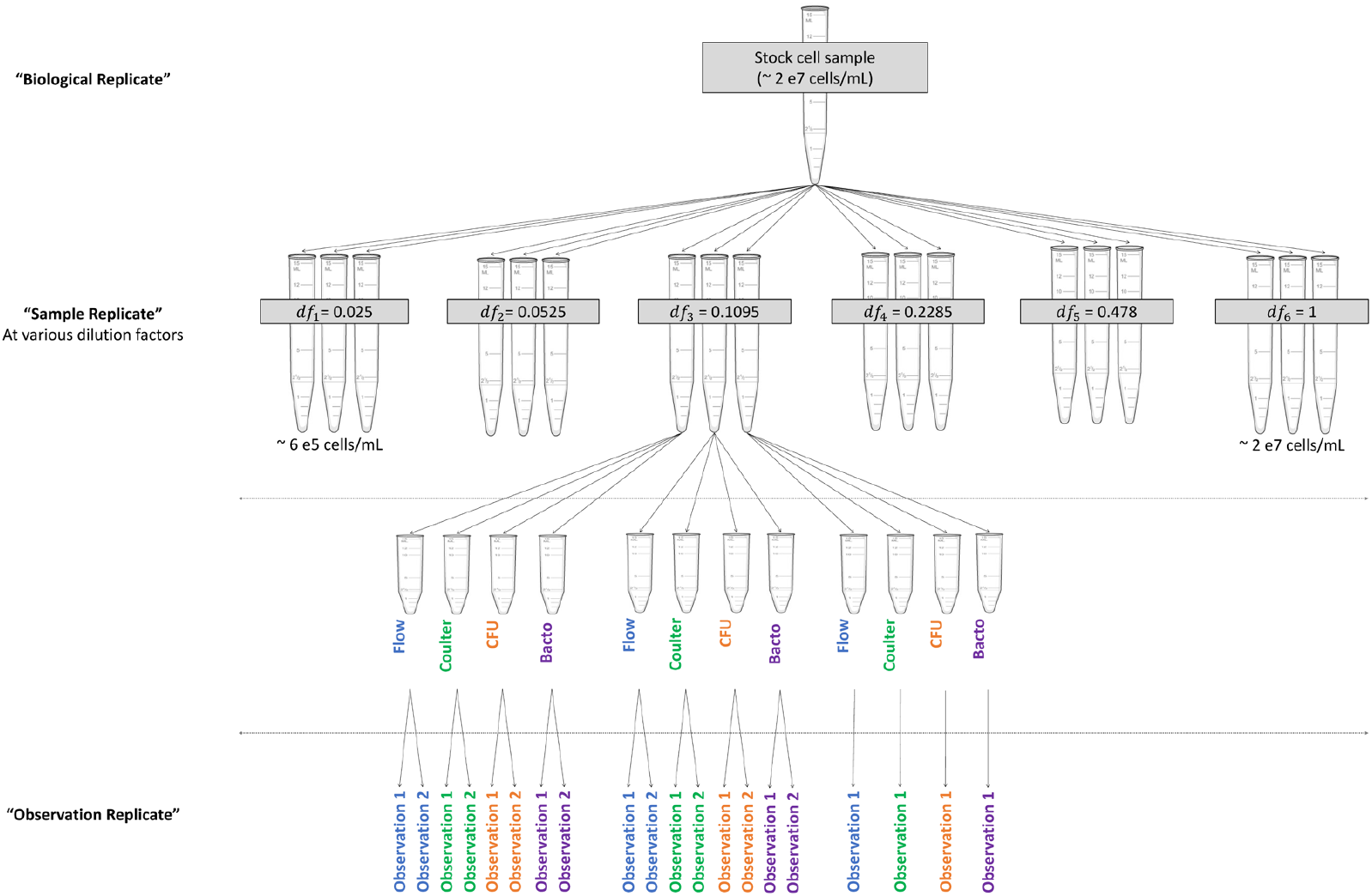
Experimental Design Diagram. Visualization of the relationships among levels of replication. The biological replicate was divided into six dilution factors (df) with three sample replicates per dilution factor. Sample replicates were divided into aliquots for each operator. Operators performed instrument-specific processing steps to record data for observation replicates.

### Bead Preparation

Bead measurements were performed on a single date using only the Multisizer and CytoFLEX. To generate the bead stock, beads (Fluoresbrite® YG Microspheres 1.00 µm, Polysciences, catalogue # 17154-10) were added to PBS to approximately 5 × 10^7^ beads/mL. This stock was diluted to eight levels evenly spaced in log-scale, and three replicate samples were prepared for each dilution factor for a total of 24 samples. The dilution factors were 1 (5 × 10^7^ beads/mL), 0.412 (2.06 × 10^7^ beads/mL), 0.1694 (8.47 × 10^6^ beads/mL), 0.0698 (3.49 × 10^6^ beads/mL), 0.0286 (1.43 × 10^6^ beads/mL), 0.0118 (5.9 × 10^5^ beads/mL), 0.00486 (2.43 × 10^5^ beads/mL), and 0.002 (1 × 10^5^ beads/mL). Samples were then divided into 2 aliquots and transferred to each operator with blinded labeling. Randomization format, observation replicate generation, and data reporting were the same as for the cell experiments.

### BactoBox

#### Data Collection

BactoBox (SBT Instruments, hardware version 7.3) was utilized per manufacturer’s protocols. PBS was diluted to 0.11X and 0.22 µm filtered before use to achieve a conductivity within the operational range of the instrument. For each replicate observation, 101 mL of sample was added to 10 mL PBS in a 15 mL conical tube, vortexed, and measured. Thus, observations are performed on concentrations approximately 100X more dilute than the prepared sample replicates. The total particle number, intact cell number, and timestamp were recorded for each sample. After each sample run, a “blank” tube was run to reduce material carryover that might otherwise artificially increase counts. For the first two biological replicates 0.11X PBS was used as a blank, and for the third biological replicate 70 % (by volume) isopropanol was used. The instrument requires disinfection with 70 % (by volume) isopropanol after every 20 measurements. When disinfection was performed, an additional 0.11X PBS blank was run afterwards to rinse the tubing.

#### Data Processing

Any count values below 10,000 were not quantifiable by BactoBox and were reported as “NA” for analysis in R. Intact counts in the blank samples were generally too low for the BactoBox to report, and thus no “background” correction was performed. Total counts were quantified for most blank samples, but no correlation was found between sample count and subsequent blank count for BioRep1 (**Supplementary Figure 1**). However, in BioRep2, the counts in the blank samples were unexpectedly high partway through the study. Therefore, the first blank after running a sample was used to estimate the “background” for each BioRep, and this was subtracted from the reported total counts. Negative values derived from this correction were converted to NA for analysis. Since BioRep4 did not use 0.11X PBS as a blank and did not have any reported blank counts, the blank values from the first two biological replicates were averaged and used for background subtraction of BioRep4. Final counts were reported after correcting for the dilution into 0.11X PBS.

#### CFU

CFU assays were performed by drip plating on Tryptic Soy Agar plates prepared from Tryptic Soy Broth (BD Bacto, catalogue # 211825) and 1.5 % (by mass) Agar (BD Bacto, catalogue # 214010). Two 96-well plates were prepared with 180 mL PBS per well. Each sample replicate was vortexed, and 20 mL of the cell sample was added to the first column in the plate to make the first 10X dilution. A multi-channel pipetter was used to serially dilute 20 mL across the next four columns (to 10^5^ dilution). For replicate observations, a second separate dilution of the sample replicate was prepared in the well plate. After completing the dilutions, the multi-channel pipetter was set to 10 mL, and dilutions 10^2^ to 10^5^ from each row were drip plated onto agar plates. Thus, observations were performed on concentrations ranging from 10^2^ times to 10^5^ times more dilute than the sample replicates. A PBS negative control plate was also prepared. Samples were incubated for 17 h to 19 h at 37 °C. Colonies were manually counted, and CFU was determined by selecting dilution(s) with colony counts between 5 to 50. If multiple dilutions with 5 to 50 colonies were present, the geometric mean was used to calculate CFU.

### Multisizer

#### Data collection

The Multisizer (Beckman Coulter Multisizer 4 with a 20 mm aperture tube) was used for all samples. First, size calibration beads (Beckman Coulter 2.0 mm Coulter CC Size Standard L2, catalogue # 6602794) were run per manufacturer’s instructions. Accuvette™ (Beckman Coulter) sample holders were prepared with 9.987 g (≈ 10 mL) + 0.05 g of 0.1 mm filtered 0.9% sodium chloride irrigation solution (Baxter, catalogue # 2F7124). Immediately before each acquisition, the sample replicate was briefly vortexed, 500 mL cells or beads were added to a prepared Accuvette, and gently inverted to mix. Thus, observations were performed on concentrations approximately 21X more dilute than the prepared samples. For replicate observations, separate Accuvettes were prepared. Samples were run with an acquisition volume of 100 mL. Between samples, the aperture tube was flushed with saline and then immersed in an Accuvette filled with saline for rinsing.

#### Data Processing

The Multisizer software was used to calculate sample concentration. The “Coincident Correction” feature was applied in the software to account for single events deemed likely to consist of two or more particles. The software automatically calculates the number of objects/mL for the original, undiluted sample replicate based on the acquired volume, dilution into saline, and selected size range. For cell runs, objects with an equivalent diameter ranging from 0.8 µm to 2.2 µm were counted. For bead runs, objects with an equivalent diameter ranging from 0.8 µm to 1.616 µm were counted.

### CytoFLEX LX

#### Data collection

Cell sample preparation: Two fluorescent probes were utilized to enable two types of count. Hoechst33342 is a cell-permeant probe that primarily binds double stranded DNA; Hoechst-positive events were classified as cells. DiBac is a cell membrane potential probe that permeates cells with a compromised membrane and Hoechst-positive/DiBac-negative events were counted as intact cells. Hoechst33342 working stock (HWS) was prepared by diluting 20 mmol/L Hoechst33342 (Thermo Fisher Scientific, catalogue # 62249) to 2 mmol/L in PBS. DiBac working stock (DWS) was prepared by diluting 2 mmol/L DiBac4(3) (Bis-(1,3-Dibutylbarbituric Acid)Trimethine Oxonol; Biotium, catalogue # 61011) in dimethylsulfoxide (DMSO) to 0.2 mmol/L in PBS. Compensation controls were generated by adding each probe to a separate tube of cells for a final concentration of 8 µmol/L Hoechst and 0.8 µmol/L DiBac. HWS and DWS were combined in equal volumes to make probe working stock (PWS). A cell-free probe control was generated from PBS and PWS at the same concentrations as the compensation controls. Controls were protected from light and incubated for at least 15 min at room temperature. After data from controls were collected, the test samples were vortexed, 500 mL was pipetted into a microcentrifuge tube, and PWS added for a final concentration of 8 µmol/L Hoechst and 0.8 µmol/L DiBac. For each replicate observation, a separate sample aliquot with probes was prepared. After probes had been added to all samples, tubes were incubated at 37 °C, 120 rpm for 30 minutes then stored at 4 °C until immediately before acquisition.

Cell and bead sample processing: Beads were used as prepared after the dilutions were generated. System startup and quality control were performed on the flow cytometer (CytoFLEX LX, Beckman Coulter) per manufacturer’s instructions. For each sample, 50 mL was diluted into 150 mL PBS in a flow tube (4X dilution). Samples were run for 4 minutes at 10 mL/min with the acquisition conditions in **Supplementary Table 1**. After each observation of a cell sample, a PBS blank was acquired for 30 s at approximately 100 mL/min to rinse. All fcs files were exported for analysis in RStudio.

#### Data processing

Separate gating schemes were designed for cell and bead samples, then applied to all relevant data files. The application of the gating scheme is demonstrated in **Supplementary Figure 2**. Final counts were reported after back-calculating the concentration based on the acquired volume and dilution used in the flow cytometry-specific sample preparation.

For cell and bead gating: The first minute of data was removed due to high rates of low scatter/Hoechst-negative events, then a logicle transform was applied to the four channels of interest (FSC, VSSC, NUV450, B525).

For the cell gating: A representative file for gate definition was generated by concatenating four files evenly spaced across an experiment’s duration, and a gate identifying cell events was defined on NUV450 and VSSC. The Hoechst-only compensation control was used to define a threshold gate in the B525 channel to separate events with high and low membrane potential (assigned to dead and viable, respectively). Data were not compensated due to the small spillover between probes (confirmed with the single-color controls) and because there were only two fluorescent channels of interest.

For the bead gating: Manual gates were added on SSC and B525 to delineate three peaks corresponding to singlet, doublet, and triplet events.

## Statistical analysis

Several metrics were computed, including proportionality index, variability, precision, and proportionality constant, to characterize and compare the behavior of each counting method. Detailed methods, including the stan model, are available in the Supplemental Information. Two outlier points were removed before analysis (**Supplementary Figure 3**).

### Proportionality Index

As outlined in ISO 20391-2:2019, an idealized counting process should follow the proportional relationship:

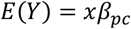

That is, the expected mean cell concentration, *E*(*Y*), is proportional to the dilution factor, *x*, with an associated *proportionality constant, β*_*pc*_. Here, *YY* is a random variable representing the observed cell concentration, and *E*(*Y*) is the expected value, or mean, of *Y*. This model implies that diluting a sample by a given factor should be reflected by a proportional decrease in the resulting measured cell concentration of that sample. Any systematic deviation from this model implies there is some discrepancy between the particular counting method and an ideal counting process. As our data are evenly spaced in log-scale, we use a similar approach, but the proportional assumption is made on the log-scale. Equally spacing the dilution factors in log-space allows for more stable statistical procedures versus applying the ISO calculations directly to the linearly scaled (raw) data, as the unequal spacing of the dilution factors on the raw scale leads to imbalances in how different observations are weighted for the various statistical methods. Furthermore, instead of analyzing concentration directly, we evaluate concentration scaled by the dilution factor, which should be a constant for a proportional relationship. The advantage of the concentration scaled by dilution factor is that it is easier to visually diagnose deviations from proportionality and differences in concentration. The proportional model can then be expressed as:

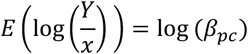

As described in the ISO 20391-2:2019 standard, we use the *Proportionality Index* (PI) as a metric of deviation from the idealized proportional model. To compute the PI, we first fit a flexible model (for example, a higher order polynomial model) that can adapt to non-proportional trends in the data. We then fit the proportional model specified above and compute a measurement of distance between the fitted flexible model and the fitted proportional model. A smaller PI indicates that the data closely follow a proportional relationship, while a larger PI suggests greater deviation from the proportional model. For this analysis we use the following PI metric based on the mean of absolute residuals for the log-scaled concentrations:

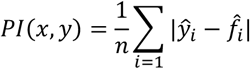

Where *ŷ*_*i*_ is the predicted log cell concentration at a log dilution factor of *x*_*i*_ for the proportional model (fitted using the data,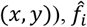 is the corresponding predicted value using the flexible model, and *n* is size of the given sample. This metric thus measures the empirical deviation between the idealized proportional model and a flexible model that is sensitive to any non-proportionality found in the observed data. The proportional model was estimated as follows:

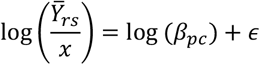

Above, *β*_*pc*_ is the true proportionality constant to be estimated, and *ϵ* is a mean zero Gaussian random variance with unknown variance *σ*^2^. Here 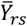 represents the average of the replicate observations in each replicate sample (following the same approach as outlined in the ISO 20391-2:2019 standard).

The flexible model is similar, but with additional terms:

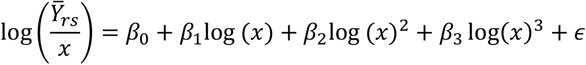

The additional components can adapt to non-proportionality in the observed data. Specifically, *β*_1_, *β*_2_, and *β*_3_represent coefficients for polynomial terms that can account for non-proportionality. Differences between the two models’ predicted mean responses indicate deviation from proportionality (with larger differences implying larger deviation from proportionality).

To compute uncertainties for the PI, we apply the non-parametric bootstrap as described in the ISO 20391-2:2019 standard. That is, we compute the PI on numerous (i.e., thousands) of resampled datasets, where each datapoint is sampled with replacement. The collection of resampled PI values constitutes an approximation of the sampling distribution of the PI, which we can use to compute confidence intervals using the appropriate quantiles from the bootstrap distribution.

### Variability and Precision

For characterizing the variability (or precision) of a cell counting method, we fit a Bayesian hierarchical model, as this approach allows us to model the nested nature of our data and to measure the various sources of variability introduced at each layer. Specifically, we account for the fact that measurements from the same biological replicate and replicate sample are not independent. Taking this structure into account allows us to estimate the magnitudes of variability at each level of the hierarchy: variability due to differences in (1) biological replicate, (2) replicate sample, and (3) measurement error of the instrument (observation replicates). We can also aggregate these sources of variability to estimate the total variability of each counting method. The hierarchical model is defined as follows:

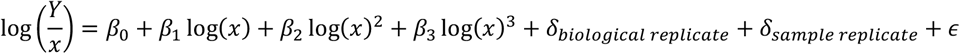

The *δ* terms above are introduced as random effects to account for the nested structure of the data. Each *δ* represents an offset from the true functional relationship to account for variability among biological replicates and between sample replicates. Each *δ* is assumed to follow a mean zero Gaussian distribution with an associated variance term for the variability due to both biological replicate and sample replicate. Uncertainties for each contribution to the overall variability are provided via the posterior distributions for the variance parameters for *δ*_*biological replicate*_, *δ*_*samplereplicate*_, and *ϵ*. The model was fit using the stan_lmer function from the rstanarm R library (Goodrich, 2024).

One consequence of this model is that it assumes that the variability of the log of the concentrations scaled by the dilution fraction is constant. Specifically, the variance at any concentration is:

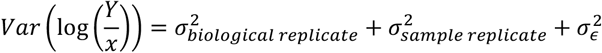

This assumption on variance holds if, on the raw scale, the standard deviation of the concentrations is proportional to the mean concentration, which is commonly observed in practice. Such an error structure implies that variability grows at a faster rate than with other common assumptions on variance, for example, the assumption that variance is proportional to the mean (as is the case with the Poisson distribution, which is a typical consideration for counting processes). If this assumption holds, then we can characterize the variability of the full dilution series at once, which allows for simple comparisons of variability between counting methods. However, if this assumption is clearly violated, then a slightly more complex model may be needed, for example, one that allows the variability in the log concentrations to change linearly with the dilution fraction.

### Proportionality Constant

The proportionality constant (PC), *β*_*pc*_, represents the magnitude of proportional change in measured concentration for a given change in the dilution factor. This value can also be interpreted as the estimated cell concentration when the stock solution from the experiment has not been diluted (i.e., dilution factor is one). The proportionality constant is estimated via fitting the proportional model to the observed data. Specifically, for each counting method, we use the following model:

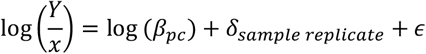

In a similar manner as above, we estimate this model using the rstanarm package, which interfaces to the stan library. We then use samples obtained from the posterior distribution *p*(*β*_*pc*_ |*x, y*) to compute estimates and uncertainties for the proportionality constant, *β*_*pc*_.

## Results

### Overview

We performed four separate experiments with cell samples and one with bead samples, and quality metrics were calculated for each dataset. We present these results as a useful case study to describe implementation of the quality metrics. Additional results are shown in **Supplementary Information**.

### Proportionality Index

A proportional model was fit to the data from each biological replicate for each method (**Figure 2A**), and the PIs were calculated (**Figure 2B**). Evaluating the proportional model fits is useful for understanding the resulting PIs, and **Figure 2A** shows fits for the separate biological replicates (solid lines). For a perfectly proportional system, the plot would show a horizontal line (dashed lines), as is close to the case for Total-A and Total-C. Total-B and Viable-B both diverge from proportionality at one end of the dilution factors, as does one replicate of Viable-D. Lastly, Viable-C has a repeatable, consistent deviation from proportionality. After understanding trends in the proportional model fit, **Figure 2B** can be easier to interpret. Total-A and Total-C have the lowest PI (indicating they are more proportional), and Viable-B and Viable-D estimates are slightly higher due to small deviations from proportionality visible in the fits. Lastly, Viable-C and Total-B have greater deviation from proportionality, though the high uncertainty in the Total-B estimates makes it more difficult to interpret.

**Figure 2.**
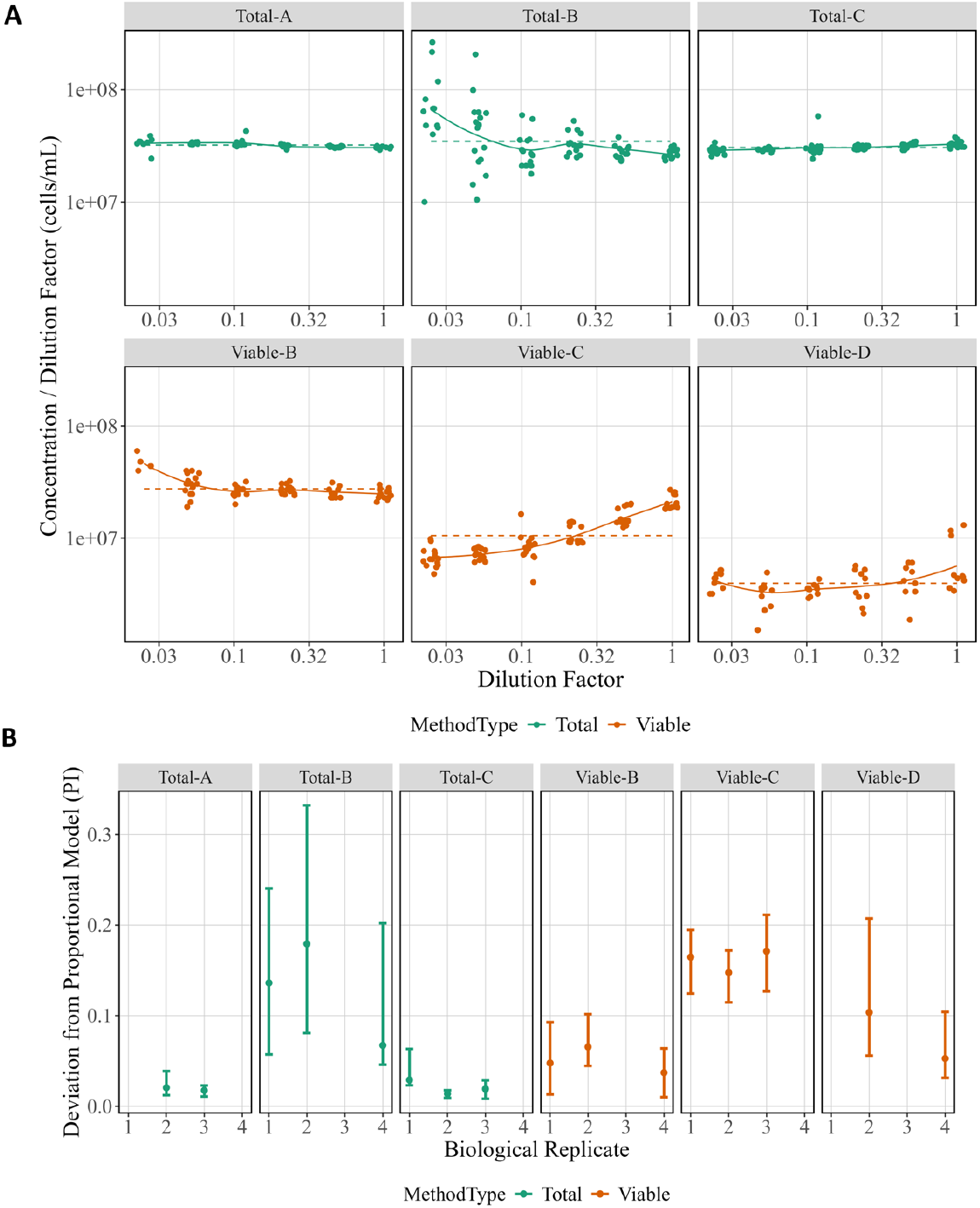
Proportional Model Fits and Proportionality Index Across Method and Biological Replicate. (A) Raw data from each biological replicate are displayed separately for each method with the y-axis giving Concentration (in cells per mL) divided by Dilution Factor and the x-axis showing Dilution Factor. Both axes are in log10 scale. Colors correspond to method type, the proportional model fit for each is indicated as a dashed line, and the non-proportional fit as a solid line. (B) The PI for each biological replicate is shown for each method. Data points represent the computed PI, and vertical bars indicate 95 % bootstrap intervals. Colors correspond to method type.

### Variability

An additional advantage of ISO 20391-2:2019-like experiments is the ability to investigate overall variability and estimate which replicate level contributed the most variability. There were large differences in overall variability for the methods tested, with all methods except Total-B having lower variability than Viable-D (CFU) (**Figure 3A**). While most methods showed constant variance across the dilution series, Total-B tended to have higher variability at the lower dilution fractions. Thus, the large amount of variability detected in the Total-B method is likely attributable to the lower half of the dilution series, and the variability may be comparable to some of the other methods if only higher dilution fractions are considered. Next, the relative sources of variability can be evaluated (**Figure 3B**). In this work, the three levels of variability are between biological replicates (days, n=2-3), sample replicates (n=18 per biological replicate), and observation replicates (n=30 per biological replicate). Total-B and Viable-D had the highest estimated variability for sample and observation replicates. Viable-C was the only method where biological replicate variability was highest. Otherwise, sample replicate variability and observation replicate variability estimates did not show distinct trends. All variabilities are given as standard uncertainties for the particular source of uncertainty. As a consequence, the total variability in Figure 3A is not a sum of the separate variabilities in Figure 3B (since, in general, the standard uncertainty of the sum of uncorrelated random components is not equal to the sum of their individual standard uncertainties).

**Figure 3:**
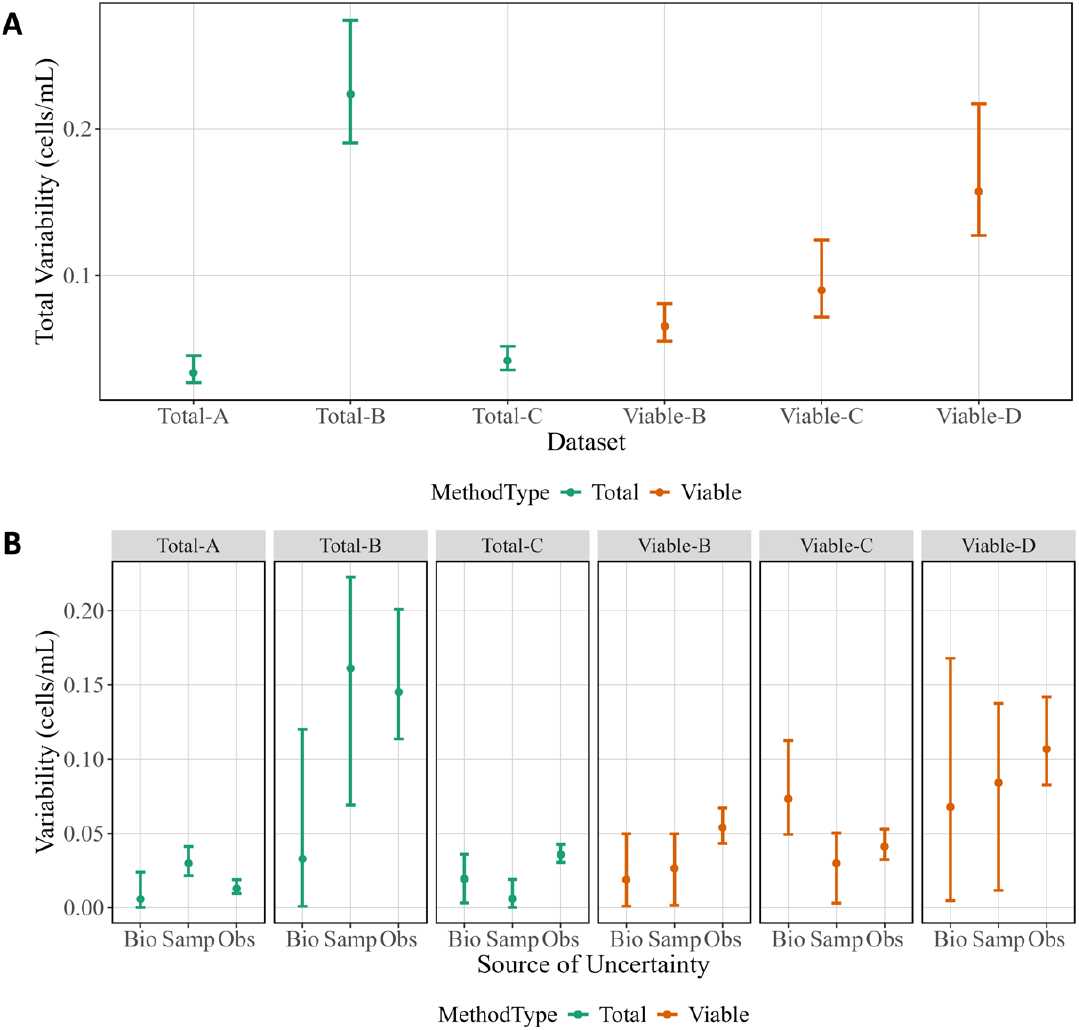
Total Variability and Sources of Variability from Cell Experiments. (A) Total variability by method. (B) Variability attributed to each replicate level by method. Data points represent posterior medians, and vertical bars indicate 95 % credible intervals. Biological replicates (Bio), sample replicates (Samp), and observation replicates (Obs) are shown for each method. Colors correspond to method types.

Variability could also come from the order in which samples were prepared and acquired, as cells could potentially die, aggregate, or multiply during the experiment. This type of order-dependent variability was minimized as the cells were kept cold in nutrient-free media during data collection. To confirm no meaningful effect was observed, the slope of the residuals was investigated as a function of the preparation order and the acquisition order. No method showed a consistent trend in residuals as a function of the acquisition order or preparation order (**Supplementary Figure 4**).

### Proportionality Constant

In addition to evaluating the quality of the measurements, the absolute estimates of cell concentration can be compared using PCs to evaluate measurement bias or true differences between methods **(Figure 4**). While the measurands differ, all of these methods are often used to report total cell or viable cell counts. Overall, there was little difference in PC estimates across biological replicates, but there were differences across methods. Total cell number and viable cell number were not expected to agree, because the starting stock was prepared from a lyophilized cell formulation known to have contain some dead cells instead of a fresh culture. The three total cell measurements have similar estimates for PC, though the Total-B estimates have larger uncertainties. Estimates from viable cell methods disagree, with Viable-B giving the highest estimate and Viable-D giving the lowest estimate. The Viable-C PC estimate, which falls between the other two, is derived from noticeably non-proportional measurements that agree more with the Viable-B measurements at high cell concentrations and Viable-D measurements at low cell concentrations (**Figure 2A**). Insights from the raw data that may explain this are detailed in the discussion section.

**Figure 4:**
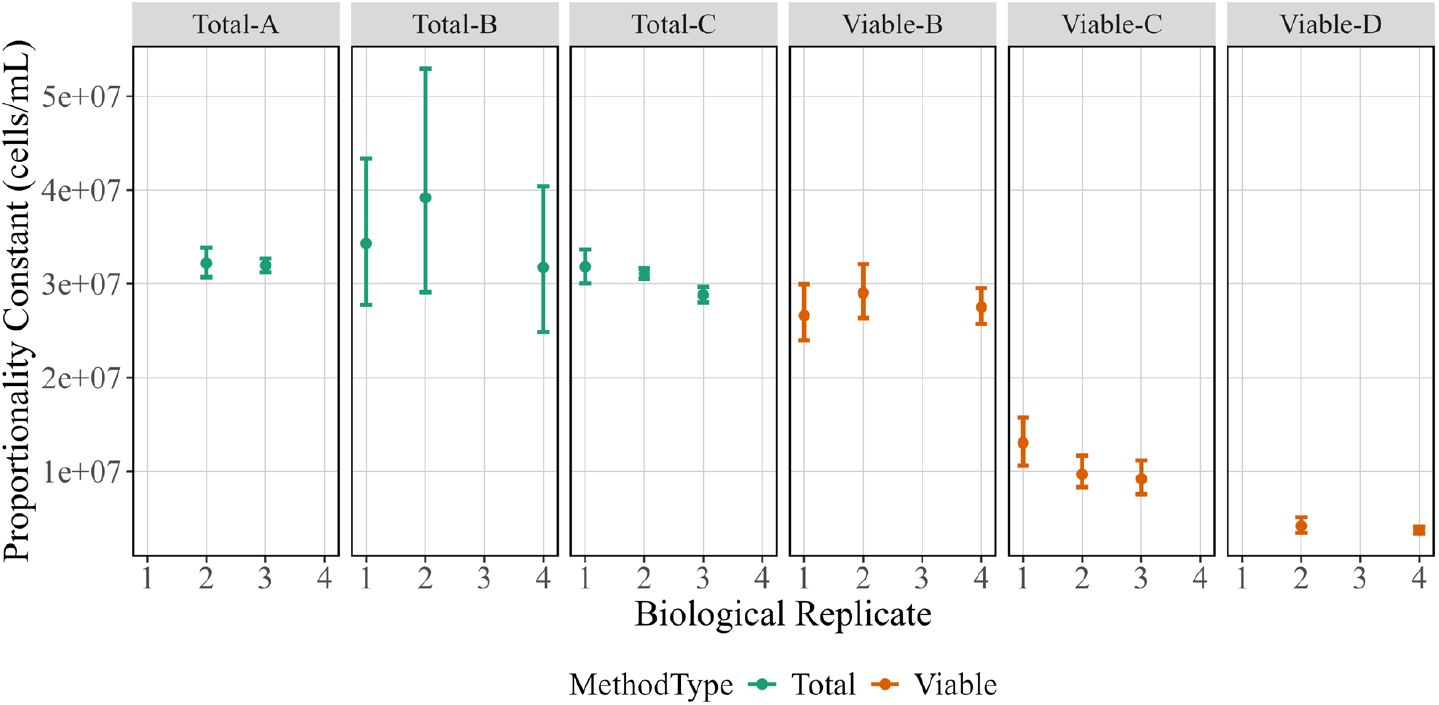
Proportionality Constants for Biological Replicates Across Method. Proportionality constant estimates (in cells/mL) are shown on the y-axis versus biological replicates are shown on the x-axis. Data points represent posterior medians and vertical bars indicate posterior 95 % credible intervals. Colors correspond to method type.

## Discussion

In this study, a new framework to assess microbial cell counting methods was developed and applied to demonstrate comparisons across common counting methods as a representative use case. The specific experimental design and data analyses are both required for the appropriate application of the framework. The ISO 20391-2:2019 experimental design and mathematical analyses for cell counting methods were modified to calculate measurement quality metrics for log-scale microbial cell datasets. Example datasets were collected to demonstrate how results and quality metrics can be used to inform fit-for-purpose method selection. Measurement quality metrics support measurement assurance and give context when results from different methods disagree. Since measurements such as total or viable cell count have no ground-truth reference materials, quality metrics are critical for evaluating relative method performance.

Proportionality over a large concentration range is particularly important for microbial cell counting methods since sample cell concentrations may span several orders of magnitude. Variation in levels of proportionality is not always noticeable on the linear scale and challenging to compare without supporting statistics, such as PI. An important caveat for PI comparisons is that the experimental design (i.e., dilution factors, sample and observation replicate numbers) must be consistent throughout all experiments to be compared. We used the PI to summarize relative proportionality (**Figure 2**). For example, Total-A and Total-C have lower PI estimates (better proportionality) than CFU (Viable-D) (**Figure 2B**), suggesting a total particle count measurement paired with CFU may be valuable for measurement assurance. Additionally, PI can be used to optimize a single method by comparing PIs from different sample processing protocols. For example, the first two Method-B experiments were performed with 0.11X PBS rinses, and the third experiment used a disinfectant rinse. It was expected that proportionality would be improved (lower PI) for whichever rinse led to greater reduction in carryover events. Although the estimated PI was lower for the third experiment, it is difficult to reach a definitive conclusion given the small sample size and large uncertainty in PI estimates for Total-B. Since proportionality is a key quality indicator for microbial cell counting methods, PI is a useful tool for intra- and inter-method comparisons.

The quality metrics can also be used to evaluate the impact of data analysis decisions on results from a single dataset. For example, the Viable-C dataset (**Supplementary Figure 5**) could be further analyzed to determine if different gating parameters result in more proportional results, or the Total-B dataset could be reanalyzed using different estimates of non-cellular “background” events. In general, thresholding leads to zero background events in blank samples for some methods, but other methods will still have measurable background events. For methods with non-zero background event counts, it is important to justify how to adjust cell counts to reflect the background since adjustments could have a significant impact on PI estimates, particularly if measurements at lower cell concentrations are of comparable magnitude to the blank. We also investigated whether our PI results would change significantly if we considered only a sub-range of the measured dilution factors (**Supplementary Figure 6**) and found only minimal changes that did not alter conclusions. However, this sub-range analysis could be valuable if sample concentrations were not in the operating range of a method, such as for Method B and the lowest cell concentration tested here. Overall, evaluating the PI is a straight-forward way to compare cell count methods.

Measurement variability is another metric that can inform method selection. The experimental design had three levels of replication (biological, sample, observation) such that variability at each level could be quantified separately, in addition to overall variability (**Figure 3**). As expected, CFU had among the highest overall variability. The analysis also breaks down the variability due to observation versus sample versus biological replicate. The goal is to use sample replicates to estimate variability due to sample preparation and observation replicates to estimate variability due to the measurement instrument or procedure. For methods where sample replicate variability is higher, more samples could be generated and measured fewer times. For methods where the observation replicate variability is higher, the opposite could be done. Evaluating the estimated variability at each replicate level therefore can help determine where to focus improvement efforts.

Another metric, the PC value, is used to estimate the starting stock cell concentration while incorporating the full dilution series via the proportional model fit (**Figure 4**). The PC values can be critical for method selection and are informational when comparing methods, because values could diverge due to measurement biases and/or differences in measurands. Some differences in PC were expected: for example, PC for total cell counts should generally be higher than PC for viable cell counts. Additionally, the three viability methods have different measurands and therefore do not assay the same cell properties. There is widespread acknowledgement that viability is more complex than a cell’s ability to multiply, and cell health is better understood as a continuum that includes metabolic activity, membrane integrity, and membrane potential (Díaz et al., 2010, Nebe-von-Caron et al., 2000). Our study emphasizes two points to consider. First, the relationship between cell health metrics may depend on sample preparation, as was seen in another method comparison study (Jordal et al., 2023). The work herein used lyophilized cells, so a portion of the cells may still have some level of membrane potential and integrity but may not be capable of growing. If a fresh culture were used, the PC values across viability methods may be in better agreement. Second, total count measurements can add crucial context to viability measurements and identify issues that might otherwise go unnoticed. For example, the Viable-C data has relatively poor proportionality that could be due to the cell sample or the counting method. We concluded it was not due to the cell sample since Viable-B measured the membrane integrity of the same samples and gave more proportional results. The non-proportional Viable-C data is also not due to the instrument since the Total-C data demonstrate that the instrument can count total cells proportionally at the given dilution factors. Therefore, we concluded that the viable cell counting protocol itself is non-proportional, likely due to the ratio of cells to available fluorescent probe, which is an important consideration when making viability measurements.

After all the analyses have been completed, results can be used to drive fit-for-purpose method selection. This selection process could place greater weight on certain analyses depending on the needs of a particular use case. For example, one possible use case is selecting a newer method to replace or support CFU, which is the gold standard for cell count but has numerous disadvantages (Jasson et al., 2010). Using the example datasets in this work, we considered whether any of the total count or viable count methods performed better than CFU (Viable-D) across all analyses. As previously stated, these conclusions are only applicable to our measurement processes and would need to be validated for other measurements or microbes. CFU was less proportional than most other methods, with Total-A, Total-C, and Viable-B methods all resulting in more desirable (lower) PI estimates. CFU was also more variable than all the methods tested here apart from Total-B. These results suggest mean that Total-A, Total-C, and Viable-B might be useful in replacing CFU measurements if they were fit-for-purpose. However, the PC values of all these methods are significantly higher than CFU, likely due to the measurands (particularly for total cell count) or measurement biases. Another option is Viable-C as that method had a PC estimate more similar to CFU, but Viable-C had a higher PI estimate which may make it a less suitable replacement. Therefore, from these specific datasets, no one method could be concluded as a perfect replacement for CFU and there would be trade-offs if a replacement needed to be selected. Other users should perform similar evaluations for their own measurement processes and measurement needs to decide which methods are fit-for-purpose. We suggest several key considerations to others wishing to apply the framework demonstrated here:

1. If possible, test the same operational range across all methods to facilitate comparisons
2. Select controls carefully if background subtraction is needed to avoid influencing PI estimates
3. Replicates on multiple days may not be critical to estimate relative method performance for some use cases
4. Log-scale lends itself to testing a wide concentration range and analyzing sub-regions as appropriate

Beyond analyses and measurands, there are other method-related considerations that may be important when determining whether a method is fit-for-purpose. Methods have a range of operator expertise requirements, associated instrumentation costs, necessary reagents, maintenance requirements, and time-to-result. For example, CFU was the least costly method to perform, but the required incubation time (overnight) increased the time-to-result as compared to other methods. The other methods report results immediately after measurement, and many fluorescent flow cytometers can operate automatically for higher throughput. Additionally, the instruments have a range of purchase costs, operator training requirements, and space requirements to consider. A fluorescent flow cytometer paired with probe selection has the greatest number of potential characterizations but requires substantially more investment into method development. The fluorescent flow cytometry dataset indicates further work investigating concentration-dependent probe partitioning into the cells or induced cell death may be needed (**Supplementary Figure 6**). All these considerations could also impact final method selection.

We emphasize again that our results apply to the *entire* measurement process, which includes the starting microbial cell material, experimental operators, experimental processing steps, instrumentation, instrument acquisition settings, and data analysis. Therefore, none of our specific conclusions are extensible to other laboratories, instrumentation, protocols, samples, or datasets. For our locked-down protocols and samples, trends were similar across biological replicates, suggesting that inclusion of multiple biological replicates in the study design may not be critical to obtain reasonable estimates of relative quality metrics in this case. Previous work counting mammalian cells using a similar approach also showed that biological replicates may not be critical in all cases to draw high-level conclusions (Pierce et al., 2023). However, results cannot be extended to any modified measurement process such as new microbes, different protocols, or even different operators; new studies would be required. A single experiment using beads was performed with two methods to see whether the observed trends would hold for a sample without biological variability. We found similar results between Bead-A and Bead-C (**Supplementary Figure 7**) which was also true for Total-A and Total-C. It is possible that some sets of microbes behave similarly enough to be modeled by a study of one microbe. For example, the results shown here might be extensible to other single cell suspensions of microbes with a similar shape, but substantially more work would be needed to evaluate this hypothesis. In other cases, new datasets may be needed to estimate the proportionality of count methods applied to other microbes. As we continue to characterize more cell counting protocols, we expect to learn more about which variables have the greatest impact on the quality metrics.

There were also limitations in the studies we executed. Sample concentrations ranged from approximately 5 × 10^5^ cells/mL to 2 × 10^7^ cells/mL; however, each operator performed additional method-specific dilutions such that the concentration range at time of measurement differed across methods. These differences were large, ranging from a 4X dilution to a 100X dilution, which resulted in some methods operating closer to an instrument’s limit of detection. Additionally, all the methods required thresholding during data analysis to classify which events are cells, intact cells, or countable colonies. Manual thresholding will introduce operator bias so automated thresholding methods can be a valuable tool for increasing analysis repeatability and transferability (Finak et al., 2016). Also, due to the large number of samples and wide range of unknown concentrations to be assayed, a drip-plate CFU method was selected, but other methods such as spread-plate CFU might have lower variability and PI. These factors are expected to have influenced our results and more studies could evaluate the impact of these limitations on the observed quality metrics.

There are many avenues to expand upon the current work such as making the experimental design and analysis easily accessible to the end user and using the demonstrated framework to improve microbial cell counting for a wide range of applications. The “COMET” RShiny application was created (Pierce et al., 2023) to help researchers implement the linear-scale analyses detailed in ISO 20391-2:2019, and a similar application could be created for the log-scale modified calculations. Future work applying the framework to improve microbial cell counting could include studies with modified protocols such that all methods measure the same range of cell concentrations to allow a more “fair” comparison of methods. The work could also be extended to other microbes of interest to determine how measurement quality metrics will vary depending on the microbe morphology or phenotype. Additionally, studies could evaluate the impact of method-specific protocol changes and use the metrics to drive protocol optimization, including data analysis optimization. Of particular interest would be enumerating multiple strains or species in a mixture. Commercial microbial products often are a mixture of microbes, and understanding how measurement quality varies across high abundance and low abundance species could help determine whether counting methods are fit-for-purpose for these mixtures. These additional studies could help inform cell count best practices for a range of samples and applications.

## Conclusion

Cell count measurements are increasingly important for many microbial industry and research applications, but CFU is by far the most common certified value found on commercial microbial reference materials. There are several reference values (CFU, membrane-intact cell number, total cell number) that would need to be certified to support all the cell counting methods tested herein. Absent such reference materials, measurement quality metrics can help support fit-for-purpose method selection. Here we introduced modifications to ISO 20391-2:2019 appropriate for microbial cell counting across log-scale concentration ranges. We demonstrated the modified experimental design and data analyses by measuring *E. coli* with three total and three viable microbial cell count methods, and discussed how measurement quality metrics can shed light on method performance. Our results showed large differences among methods in terms of the estimated proportionality index, variability, and proportionality constant, all of which should be considered when determining whether a method is fit for a particular purpose. Total particle measurements from different methods resulted in approximately the same estimated concentration, but with method-specific PI estimates and variability. The total count methods tested here relied on three different measurands, so it is notable that the PC values are relatively close. Similarities between orthogonal methods such as these are encouraging when working towards a ground-truth value. Viable count estimates were more divergent between methods due to differences in measurands and likely measurement bias as well, emphasizing the importance of multi-method characterizations. Other microbial researchers can apply the experimental design and data analyses to evaluate their samples, methods, and protocols to select fit-for-purpose measurement processes and increase measurement assurance for microbial cell counting in a wide range of applications.

### Disclaimer

Certain commercial materials and equipment are identified to specify the experimental procedure. In no instance does such identification imply recommendation or endorsement by NIST or that the material or equipment identified is necessarily the best available for the purpose

## Data accessibility

Raw data and analysis code are available: https://doi.org/10.18434/mds2-3410

## Footnotes

1 Certain commercial materials and equipment are identified to specify the experimental procedure. In no instance does such identification imply recommendation or endorsement by NIST or that the material or equipment identified is necessarily the best available for the purpose

