## Supplementary Information for "Measurement Quality Metrics to Improve Absolute Microbial Cell Counting"

**Disclaimer**: Certain commercial materials and equipment are identified to specify the experimental procedure. In no instance does such identification imply recommendation or endorsement by NIST or that the material or equipment identified is necessarily the best available for the purpose

**Data accessibility**: Raw data and analysis code are available: https://doi.org/10.18434/mds2-3410

**BactoBox Events in Blank Samples**

Blank samples (0.11X PBS) were measured after each cell sample to determine the degree of carryover between samples (**Supplementary Figure 1**). It was expected that more concentrated cell samples (higher dilution factors) would result in more carryover events (higher counts in the blanks). However, data from BioRep1 show there was no observed correlation between the dilution factor and the blank sample counts. This finding suggests that background events or carryover may potentially be due to the PBS diluent or lyophilization matrix present in the sample.

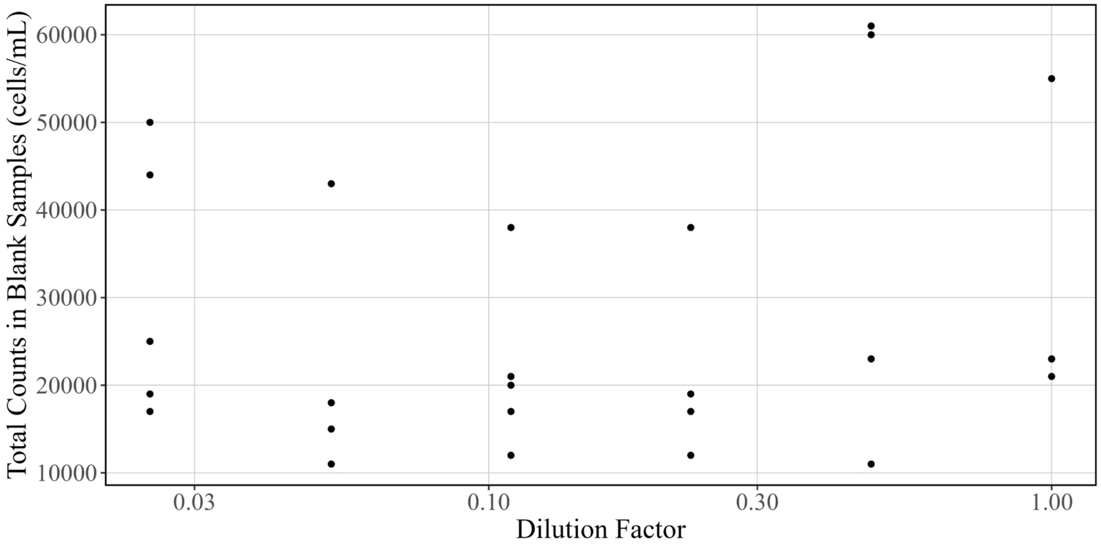


**Supplementary Figure 1: BactoBox Events in Blank Samples**. After each cell sample, a "blank" sample (0.11X PBS) was recorded. Total counts from those samples are on the y-axis, and the dilution factor of the preceding cell sample is given on the x-axis (in log-scale). Each dilution factor had five total replicates but values that measured below the BactoBox quantification limit were excluded.

### Flow Cytometry Methods

Additional details are provided for the CytoFLEX settings used during the experiment (**Supplementary Table 1**) and the gating scheme (**Supplementary Figure 2**). The full gating scheme (as an RMarkdown document) is available alongside the raw data.

**Supplementary Table 1: CytoFLEX Instrument Settings**. List of settings used for the cell or bead experiments. Letter and number combinations are listed as used by the manufacturer to specify channels.

| Setting | Cells | Beads |
|---|---|---|
| Threshold | VSSC-Height = 4000 | FSC-Height = 3000 |
| Width | VSSC | FSC |
| FSC Gain | 1000 | 100 |
| VSSC Gain | 1 | 1 |
| NUV450 Gain | 50 | 25 |
| B525 Gain | 100 | 50 |

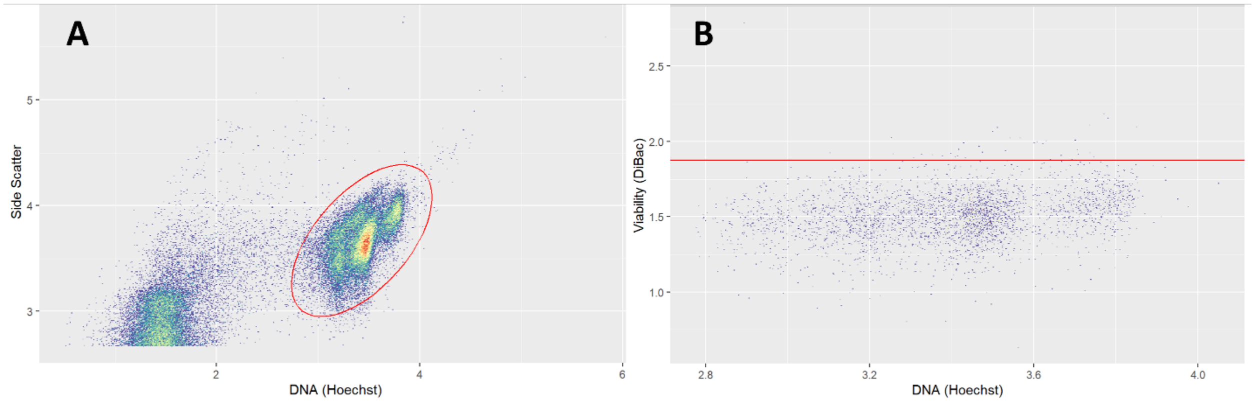


**Supplementary Figure 2: Fluorescence Flow Cytometry Gates.** Demonstration of the fluorescence flow cytometry gating scheme applied to representative data from BioRep2. All axes are shown after logicle transformation and labeled with the purpose and probe name. (A) Gate used to determine total cells shown on a concatenated file derived from samples taken throughout the experiment duration. All events within the gate (red oval) were counted. (B) Gate used to determine viable cells, shown on the Hoechst-only stained control. All events with intensities lower than the threshold (red horizontal line) were classified as live cells.

### Statistical Modeling

Statistical inference was carried out on the proportional model and flexible models in the Bayesian framework via MCMC using the rstan R library interface to the stan software library. The stan files for the proportional and flexible models are available at: https://doi.org/10.18434/mds2-3410. Half-normal priors were used for the hierarchical variance terms with standard deviation equal to the total variability seen in the data. This weighting puts a relatively large amount of prior density over values similar to those seen in the data, with relatively little mass put on much higher magnitudes. This choice of prior is considered weakly informative (and is a common default), as the resulting inference is dominated by the observed data while ruling out parameter values that are substantially higher than those suggested by the data.

### Outliers

Two data points were identified as outliers and removed from analysis (**Supplementary Figure 3**).

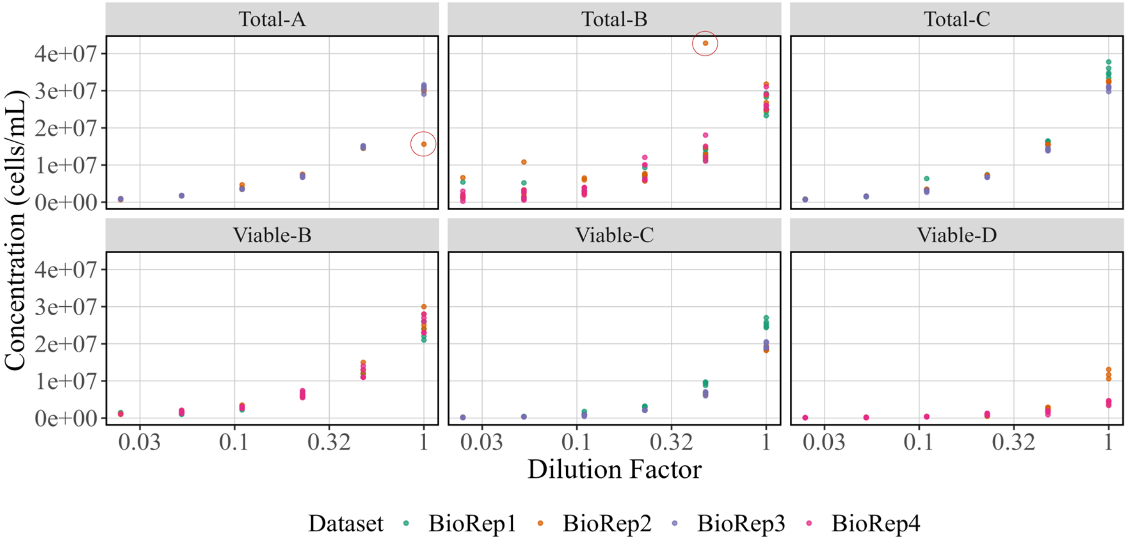


**Supplementary Figure 3**: **Outliers Identified.** Raw data are shown for each method with dilution factors on the x-axis and measured cell concentration (in cells/mL) on the y-axis. Colors correspond to biological replicates. Outliers removed from the datasets before analysis are shown with red circles.

### Acquisition Order and Preparation Order

It is possible that the order of the samples could impact the results, for example, by introducing sample carryover or by viability changing over time. During the experiment, samples were prepared by one operator from most to least concentrated, and then all instrument operators acquired the samples using a fixed sample order that included randomization. The BactoBox and CytoFLEX protocols included rinse steps between each observation as instrument documentation suggested a carryover rate of 2 % to 3 % and 1 %, respectively.

The two potential sample orders investigated were (1) the order that replicate samples were prepared from the stock solution and (2) the order that measurements were collected. To evaluate the potential effect of each sample order, residuals from the proportional model fit were evaluated as a function of order to confirm no time-dependent trends in the data (**Supplementary Figure 4**). Greater variability is seen in Total-B as a function of preparation order, but this is likely because preparation order directly correlated with the dilution factor. Samples measured in Total-B were the most dilute overall due to the method-specific dilution used and more dilute samples are expected to have greater variability in results.

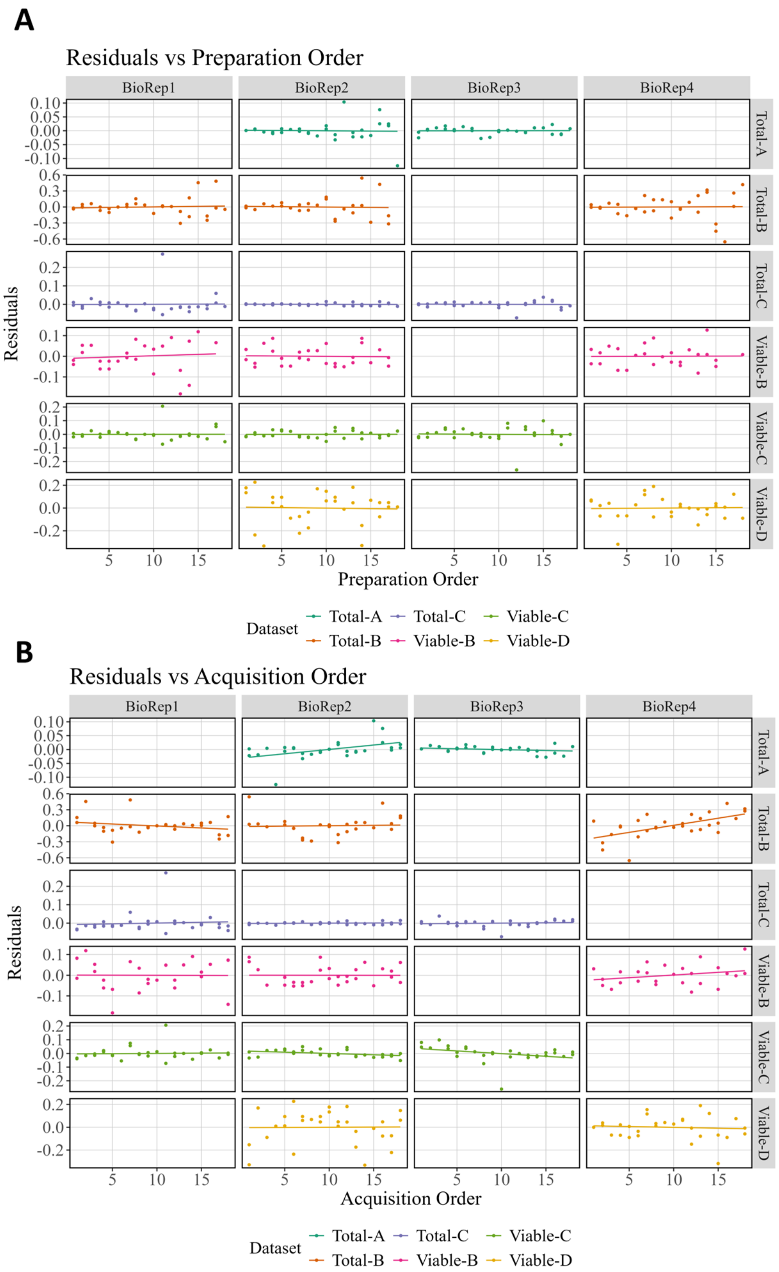


**Supplementary Figure 4: Evaluation of Time-Dependence in Results**. Residuals from the model fit were plotted as a function of the two different orders to evaluate potential time dependence. (A) Residuals as a function of sample preparation order show no consistent trends. (B) Residuals as a function of sample acquisition order show no consistent trends.

**Fluorescence Flow Cytometry Raw Data**

Raw fluorescence flow cytometry data plots of NUV450 (Hoechst 33342, all cells) and B525 (DiBac4(3), presumed dead cells) show that the distributions of B525 intensities differ depending on the underlying cell concentration (**Supplementary Figure 5**). While the threshold gate is useful to classify "viable" and "non-viable" events, the full distributions show that the data do not have distinct high and low populations in B525. There are many potential explanations for this trend. The simplest explanation would be that there is not enough probe in the solution to saturate the higher cell concentrations; in that case, increasing the concentration of probe should result in the same B525 distribution at all dilution factors. However, the concentration used was quite high. An alternative explanation is that the cells are saturated with probe even at the highest cell concentration, and that the change in B525 distribution results from cells dying as the ratio of probe-to-cell increases. These potential explanations will need further testing.

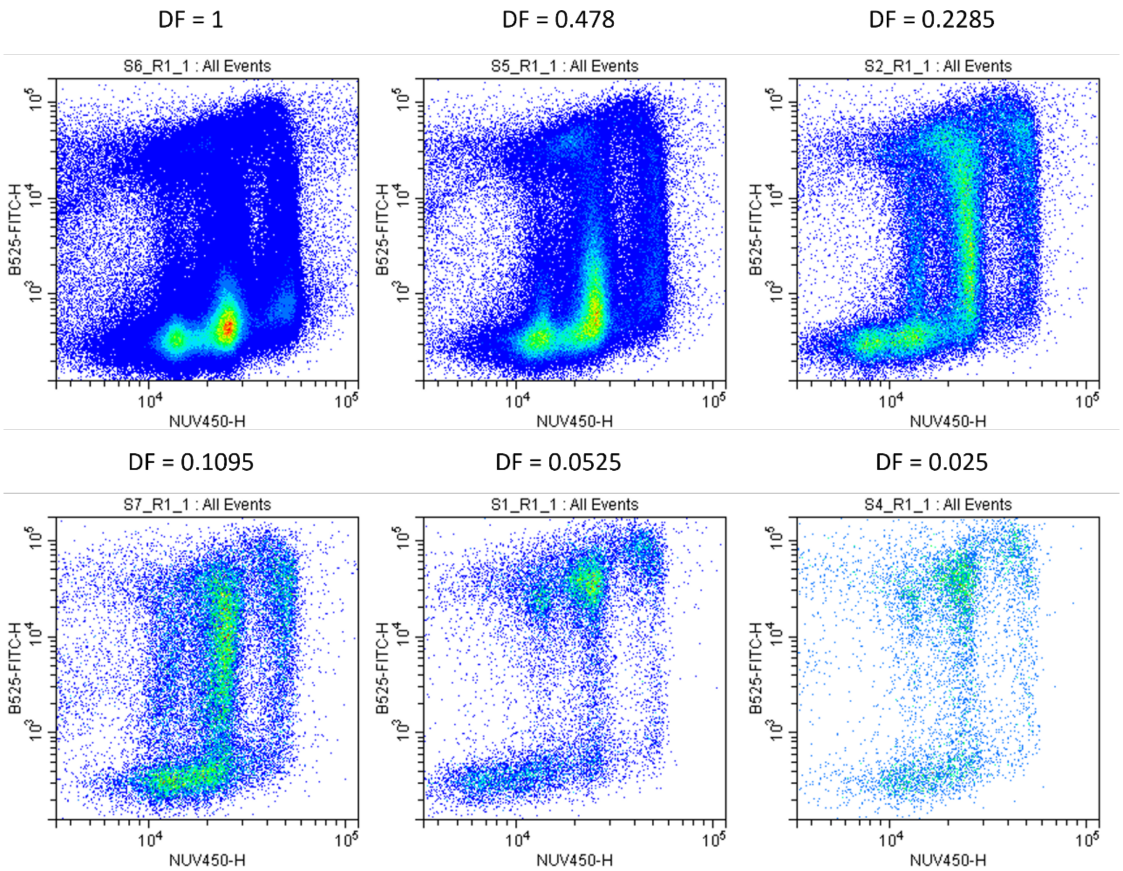


**Supplementary Figure 5: Trends in Fluorescence Intensity Distributions.** Representative fluorescence flow cytometry data from BioRep2 for samples at each dilution factor (DF) were compared. For each plot, the x-axis represents the measured intensity in the NUV450 channel, and the y-axis represents the B525 channel, and datapoints are color-coded with sparse areas of the plot in blue or green and areas with higher event density in red or yellow. The density color-coding is on the same absolute scale across all plots, indicating that the DF=1 sample has the highest cell concentration overall.

**Proportional Sub-ranges**

Our main analysis used data from all dilution factors, but measurements from some high or low dilution factors appeared to be less proportional.  For some methods, reducing the range of analyzed dilution factors resulted in lower estimates of deviation from proportionality (**Supplementary Figure 6**). For example, removing the lowest dilution factor reduces the PI of Total-B and Viable-B and better represents the manufacturer's recommended concentration range. However, all methods must be compared across the same set of dilution factors and the ranking of PI values changes little when the lowest dilution factor is excluded (**Supplementary Figure 6**, right panel). Notably, it may be difficult to derive actionable data from sub-range studies because they depend on dilution factors and the unknown starting concentration. Proportionality is likely dependent on the absolute cell concentration and the instrument's limit of detection. Here, the different method-specific dilutions mean that the measured cell concentrations differ between methods, so sub-range comparisons might not be appropriate and could be misleading.

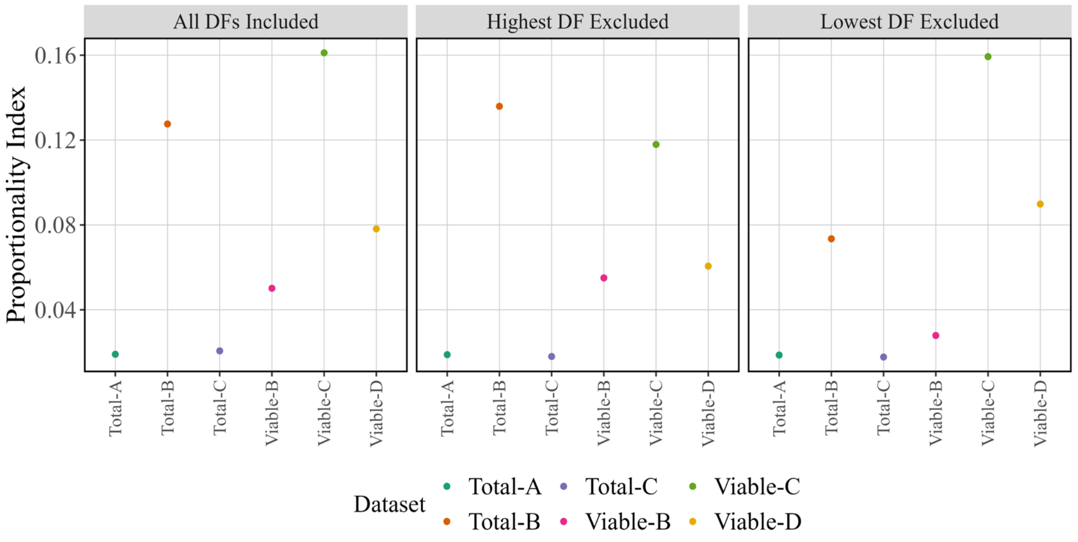


**Supplementary Figure 6**: **Proportionality Index across Dilution Factor Sub-ranges**. The average PI estimate over the entire range of dilution factors (left) was compared to the average PI computed after removing the highest DF (middle) or lowest DF (right). The y-axis shows PI, and the x-axis gives the different methods.

**Bead Experiment Overview**

A bead-based counting experiment was performed for two of the total count methods and analyzed with the same the statistical analysis carried out on the cell data. Below we provide results for the PI, variability, and PC. Overall, we conclude that PI and variability for the beads and microbial cells are similar for the tested methods, suggesting there is no large impact of the microbial cell biology on total counting performance for these methods.

**Proportionality Index**

For the bead experiment, Bead-C was more proportional than Bead-A due to the noticeable deviation in the Bead-A raw data (**Supplementary Figure 7**). This difference may be because the bead experiment covered a wider range of dilution factors (from 0.002 to 1), and the two instruments may have different limits of detection.

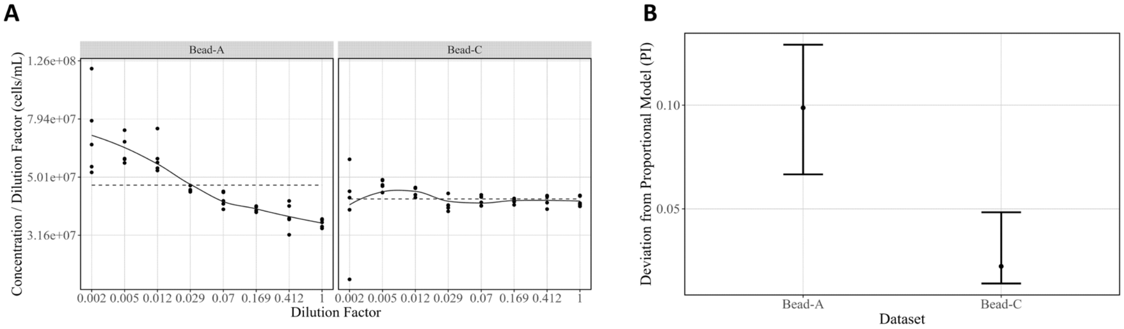


**Supplementary Figure 7: Proportionality Index for Bead Data**. (A) Proportional model fit (dashed line) to raw data. (B) Proportionality indices across methods (points represent posterior medians and vertical bars represent a 95 % confidence interval computed from the bootstrap distribution).

**Variability**

For the bead experiment, both methods showed similar total variability, and these variabilities were comparable to the results from the cell data (**Supplementary Figure 8A, Figure 3**). Additionally, the variability by replicate level appeared similar across methods, though variability at the observation level as opposed to the sample level was slightly higher for Bead-A (**Supplementary Figure 8B**).

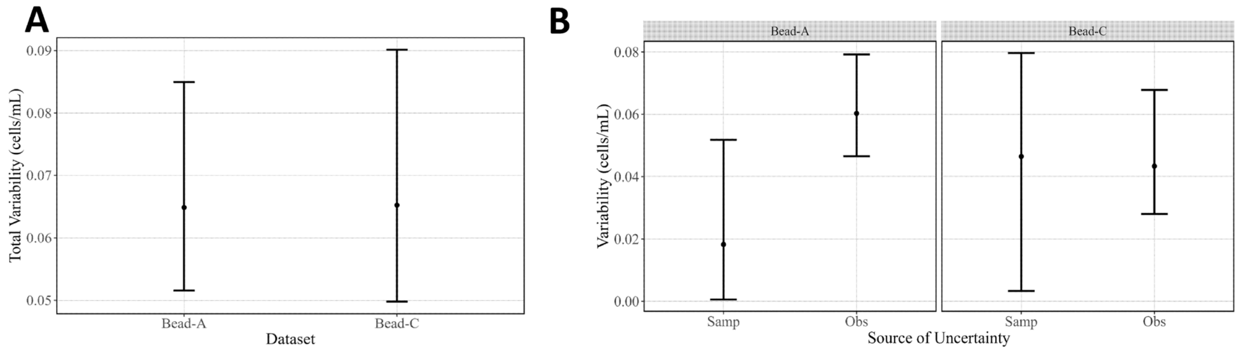


**Supplementary Figure 8**: **Variability in Bead Data**. (A) Total variability by method. (B) Variability attributed to each replicate level by method. Data points indicate posterior medians, and vertical bars represent posterior 95 % credible intervals.

#### Proportionality Constant

The two bead methods resulted in similar values for the PC (**Supplementary Figure 9**), though the median value for Bead-A was higher than for Bead-C. This trend was similar to the cell datasets where the Total-A and Total-C measurements gave non-actionable differences in PC estimates. The PC value estimates the starting stock concentration and therefore is not expected to be the same for bead and cell data.

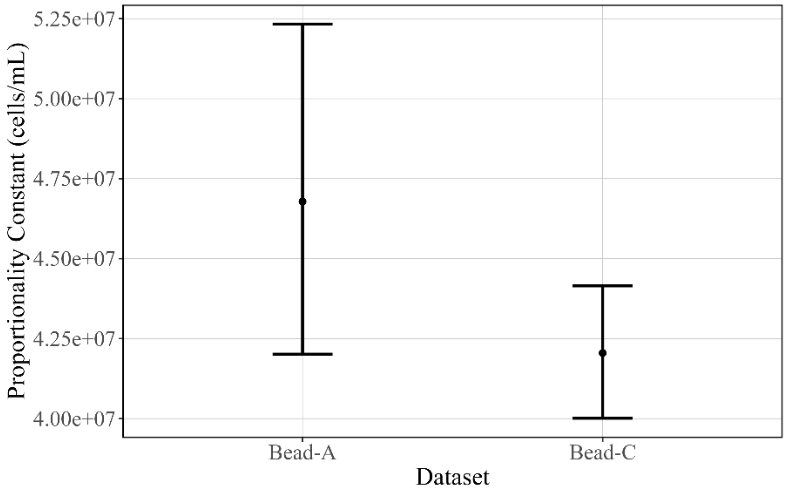


**Supplementary Figure 9**: **Proportionality Constants for Bead Data**. Proportionality constant estimate derived from each method. Points represent posterior medians, and vertical bars indicate the posterior 95 % credible intervals.
